# Secreted microbial metabolites modulate gut immunity and inflammatory tone

**DOI:** 10.1101/2019.12.16.861872

**Authors:** Rabina Giri, Emily C. Hoedt, Khushi Shamsunnahar, Michael A. McGuckin, Mark Morrison, Robert J. Capon, Jakob Begun, Páraic Ó Cuív

**Author notes:** Equal contribution (Jakob Begun, Páraic Ó Cuív). Corresponding authors: Jakob Begun (Immunology,) & Páraic Ó Cuív (Microbiology,). **ECH**, APC Microbiome Institute & Department of Microbiology, National University of Ireland, Cork, Ireland; **MMcG**, The University of Melbourne, Victoria, Australia; **PÓC**, Microba Life Sciences Ltd, Queensland, Australia.

## Abstract

Evidence is emerging that microbiome–immune system crosstalk regulates the tenor of host intestinal immunity and predisposition to inflammatory bowel disease (IBD). We identified five NF-κB suppressive strains affiliated with *Clostridium* clusters IV, XIVa and XV that independently suppressed secretion of the chemokine IL-8 by peripheral blood mononuclear cells and gut epithelial organoids from healthy human subjects, as well as patients with the predominant IBD subtypes, Crohn’s disease and ulcerative colitis. The NF-κB suppressive *Clostridium bolteae* AHG0001, but not *C. bolteae* BAA-613, suppressed cytokine-driven inflammatory responses and endoplasmic reticulum stress in gut epithelial organoids derived from *Winnie* mice that develop spontaneous colitis. This predicted *in vivo* responses thereby validating a precision medicine approach to treat *Winnie* colitis and suggesting the microbiome may function as an extrinsic regulator of host immunity. Finally, we identified a novel molecule associated with NF-κB suppression indicating gut bacteria could be harnessed to develop new therapeutics.

## Introduction

The human gut is the largest immune organ of the body and gut epithelial cells play a key role in the establishment and maintenance of gut homeostasis, as well as rapid responses to infection^1^. The gut is colonised by a diverse microbiota that has co-evolved with its host and forms a durable symbiotic relationship through its modulation of innate and adaptive immune responses^2, 3^. However, with a few notable exceptions^4, 5^ the microbes and microbial determinants of immune tone remain cryptic.

Inflammatory bowel disease (IBD) is comprised of two predominant subtypes, termed Crohn’s disease (CD) and ulcerative colitis (UC), that are characterised by relapsing and remitting gut inflammation. The Nuclear factor-κB (NF-κB) family of transcription factors are master regulators of gut epithelial integrity and inflammation, activation of antigen presenting cells and effector leukocytes, and are important contributors to the pathogenesis of IBD. Upon activation, NF-κB dimers translocate to the nucleus where they regulate transcription of a wide range of genes including those involved in immune and inflammatory responses^6^. In the healthy gut, NF-κB activation is tightly regulated^7^ however several IBD genetic risk alleles including *nod2, TOLLIP* and *A20* exert their pathogenic effects at least in part through dysregulated NF-κB signalling^8^. Additionally, CD disease phenotype correlates with NF-κB activation^9^, and macrophages and epithelial cells isolated from inflamed intestine of CD and UC subjects show increased activation of nuclear NF-κB-p65^10^. As such, NF-κB signalling contributes significantly to the cascade of host-responses underlying the pathogenesis of IBD.

The gut microbiota is increasingly recognised as an important contributory risk factor for IBD. Underlying this, the healthy and IBD gut microbiota differ and are characterised by structure-function alterations to the microbiota^11 12^, and faecal transplantation has proven effective in some patients with UC^13, 14^. Such findings suggest that key members of the microbiota regulate host inflammatory responses. Indeed, several bacterial taxa are not only more abundant in the healthy gut but can also suppress inflammatory responses and alleviate inflammation in animal models of disease^15–17^. These “anti-inflammatory” properties are best characterised for the gut bacterium *Faecalibacterium prausnitzii* A2-165 which produces secreted peptides derived from the Mam protein that suppress NF-κB in human gut epithelial cells and murine colitis^18^. However, while Firmicutes-affiliated Clostridia are amongst the most abundant and functionally diverse gut bacteria, Mam is largely restricted to members of *Faecalibacterium* spp., and much remains to be discovered about the immunomodulatory capacities inherent to other Firmicutes.

Here, we identified five new Firmicutes isolates that are comparable to *F. prausnitzii* A2-165 in their NF-κB suppressive potency, and whose activities are characterised by strain specific differences. Notably, these bacteria suppressed cytokine mediated IL-8 secretion in CD and UC derived organoid cultures and peripheral blood mononuclear cells (PBMCs). Based on these observations, we demonstrated using two *Clostridium bolteae* strains how a “precision medicine” approach can be used to predict immunomodulatory and mucosal healing bioactivity *in vivo* using the *Winnie* murine model of spontaneous colitis, demonstrating the potential of bioprospecting the human microbiome for therapeutic leads.

## Results

### Gut clostridia can suppress NF-κB

We assessed the NF-κB suppressive capacity of cell free supernatants (CS) derived from 23 Firmicutes affiliated gut bacteria previously isolated by us via metaparental mating from a healthy pre-adolescent child^19^. The isolates were principally affiliated with *Clostridium* cluster XIVa, with several isolates also affiliated with clusters IV, XV and XVIII. The isolates are distantly related to *F. prausnitzii* A2-165 and another NF-κB suppressive bacterium, *Enterococcus faecalis* AHG0090, that was also isolated by metaparental mating^20^ (Figure 1A).

**Figure 1.**
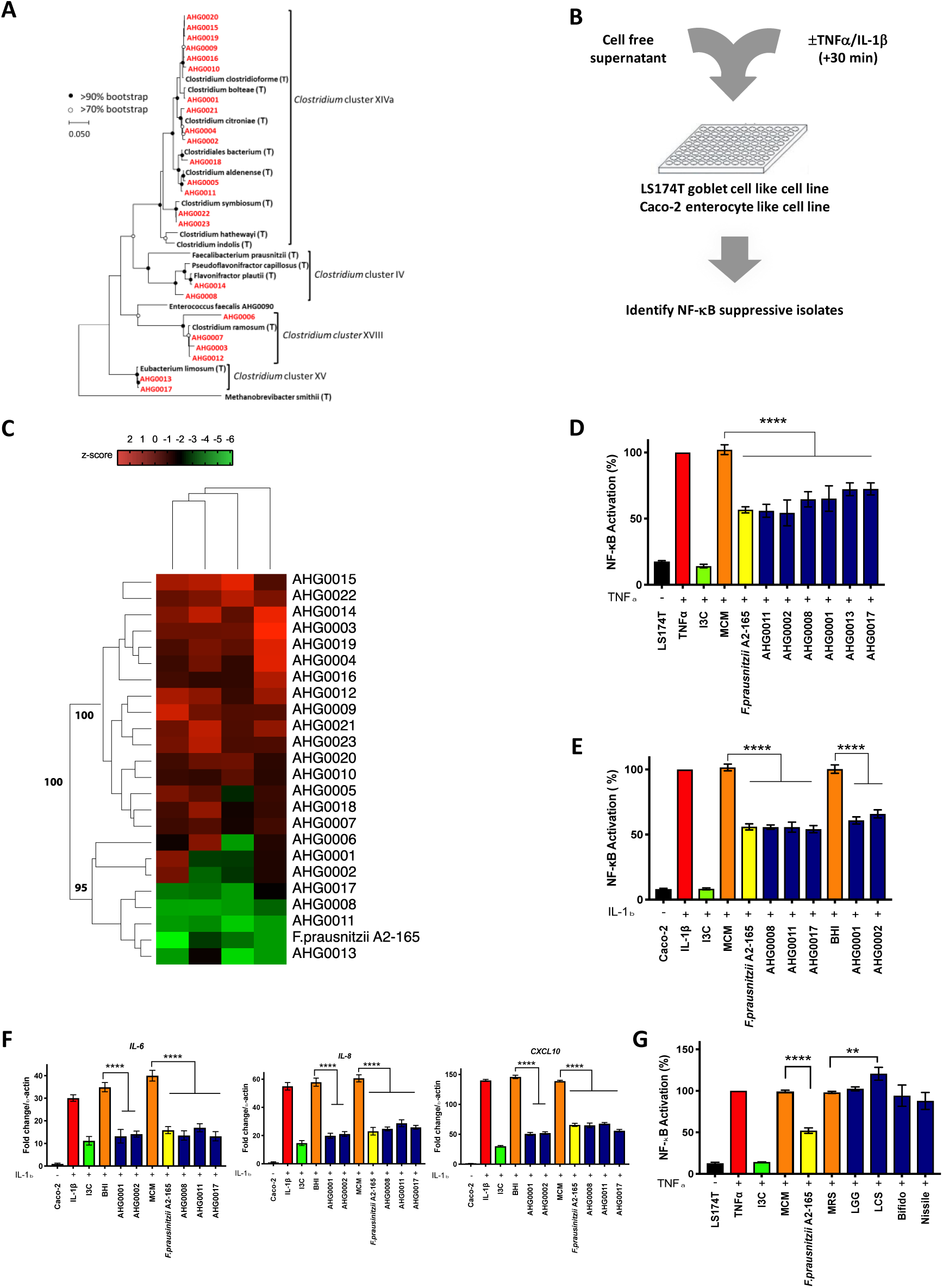
**A.** 16S rRNA based phylogeny of the MPM isolates characterised in this study (red typeface) and representative microbial isolates and reference sequences (bold black typeface). **B.** An overview of the experimental approach to characterising the NF κB suppressive capacity of the bacteria examined in this study. Cell free CS was added to the reporter cell lines which were then stimulated with cytokine after 30 min. Luciferase activity was assayed after 4 hours. **C.** Heat map analysis of the NF-κB suppressive capacity of the bacterial isolates. The ability of CS prepared from bacterial isolates grown in MCM or BHI to suppress NF-κB in LS174T or Caco-2 reporter cell lines was assessed twice independently. A Z-factor of 0.805 ± 0.06 (MCM) and 0.87 ± 0.01 (BHI) was achieved for the LS174T cells while a Z-factor of 0.78 ± 0.057 (MCM) and 0.765 ± 0.02 (BHI) was achieved for the Caco-2 cells. A subset of strains formed an NF-κB suppressive cluster with *F. prausnitzii* A2-165. **D.** LS174T based confirmatory assay of the hits identified from the first pass screen. NF-κB activation was assessed 4 h after TNFα stimulation and the extent of suppression was assessed against sterile medium (mean (standard deviation (SD))). **E.** Caco-2 based confirmatory assay of the hits identified from the first pass screen. NF-κB activation was assessed 4 h after IL-1β stimulation and the extent of suppression was assessed against sterile medium (mean (SD)). **F.** Caco-2 based qRT-PCR confirmatory assay of the hits identified from the first pass screen (mean (SD)). *F. prausnitzii* A2-165 and the validated hits suppress IL-1β induced *cxcl10*, *il6* and *il8* expression in Caco-2 cells. **G.** Analysis of the ability of *L. rhamnosus* GG (LGG), *L. casei* Shirota (LCS), *E. coli* Nissle 1917 (Nissle) and *B. animalis* subsp. *lactis* BB-12 (Bifido) to suppress TNFα mediated activation of NF-κB in LS174T cells. Cell free CS prepared from these probiotic strains do not suppress NF-κB activation in the LS174T cell line (mean (SD)). ** *p*<0.01, **** *p*<0.0001 as determined by one-way ANOVA with Dunnett’s multiple comparison test.

We assessed the ability of individual isolates to suppress NF-κB activation using LS174T goblet cell-like and Caco-2 enterocyte like reporter cell lines^21^. The LS174T and Caco-2 cells carry an NF-κB inducible luciferase reporter gene and are responsive to TNFα and IL-1β stimulation, respectively^20, 21^. Although short chain fatty acids are posited to supress gut inflammation, similar to previous reports^22^, the addition of up to 16 mM of the short chain fatty acids acetate, butyrate and propionate did not activate NF-κB under basal conditions. However, all three short chain fatty acids enhanced cytokine-driven NF-κB activation in a largely dose dependent manner (Supplementary Figure 1A-F). CS prepared from isolates following growth in Modified Clostridial Medium (MCM) or Brain Heart Infusion (BHI) medium were assessed for their ability to suppress NF-κB (Figure 1B). As previously observed^20, 21^, there was a high degree of concordance between the LS174T and Caco-2 reporter cell lines with 7 strains identified that exhibited potent activities similar to *F. prausnitzii* A2-165 (Figure 1C, Z score ≤-3). In addition to *F. prausnitzii* A2-165, the isolates *C. bolteae* AHG0001, *Clostridium citroniae* AHG0002 *Pseudoflavonifractor* sp. AHG0008, *Clostridium aldenense* AHG0011, *Eubacterium limosum* AHG0013 and *E. limosum* AHG0017 suppressed NF-κB in both cell lines when grown in MCM and/or BHI medium (Figure 1C, Z score ≤-3). The first pass screen was confirmed with CS prepared from individual isolates following growth in MCM or BHI suppressing NF-κB activation in both cell lines (Figure 1D-E, *p*<0.0001). As anticipated, the NF-κB inhibitor indole-3-carbinol (I3C) and *F. prausnitzii* A2-165 suppressed cytokine-driven activation of the luciferase reporter in the LS174T and Caco-2 cell lines (Figure 1D-E, *p*<0.0001). Consistent with the reporter assay results, all the isolates suppressed induction of the NF-κB regulated genes *mcp-1*, *il-6* and *il-8* in Caco-2 (Figure 1F) and LS174T (Supplementary Figure 1G) cells following stimulation. Critically, none of the CS exhibited cytotoxic effects (Supplementary Figure 1H-I).

We also examined the NF-κB suppressive activity of CS prepared from the widely used probiotic strains *Lactobacillus rhamnosus* GG (LGG), *Lactobacillus casei* Shirota (LCS), *Escherichia coli* Nissle 1917 (Nissle) and *Bifidobacterium animalis* subsp. *lactis* BB-12 (Bifido). Notably, none of these strains suppressed cytokine driven NF-κB activation (Figure 1G) potentially explaining the limited efficacy of probiotics for the treatment of IBD^23–27^.

### NF-κB suppression is strain specific

Having confirmed their suppressive activity, we next examined the intraspecies variations in NF-κB suppressive capacity. Isolates *C. bolteae* AHG0001 and ATCC BAA-613 (OTU1), *C. citroniae* AHG0002 and AHG0004 (OTU2), and *C. aldenense* AHG0011 and AHG0005 (OTU3) are assigned to the same operational taxonomic units (Figure 1A, ≥97% 16S rRNA sequence identity). However, these OTUs were characterised by marked intraspecies differences in their NF-κB suppressive capacities (Figure 2A-C). Next, we examined the apparent effect of growth medium on the suppressive effects of *C. bolteae* AHG0001 and *C. citroniae* AHG0002 in the first pass screen. We determined that CS prepared from *C. bolteae* AHG0001 and *C. citroniae* AHG0002 following growth in MCM but not BHI suppressed TNFα-driven mediated NF-κB activation in LS174T cells (Figure 2D). Conversely, CS prepared from these strains following growth in BHI but not MCM suppressed IL-1β-driven NF-κB activation in Caco-2 cells (Figure 2E). Thus, NF-κB suppressive functionality is strain specific and nutritional influences on bioactive production may affect the production of suppressive activity *in vitro* and the extent of anti-inflammatory activity *in situ* in the gut.

**Figure 2.**
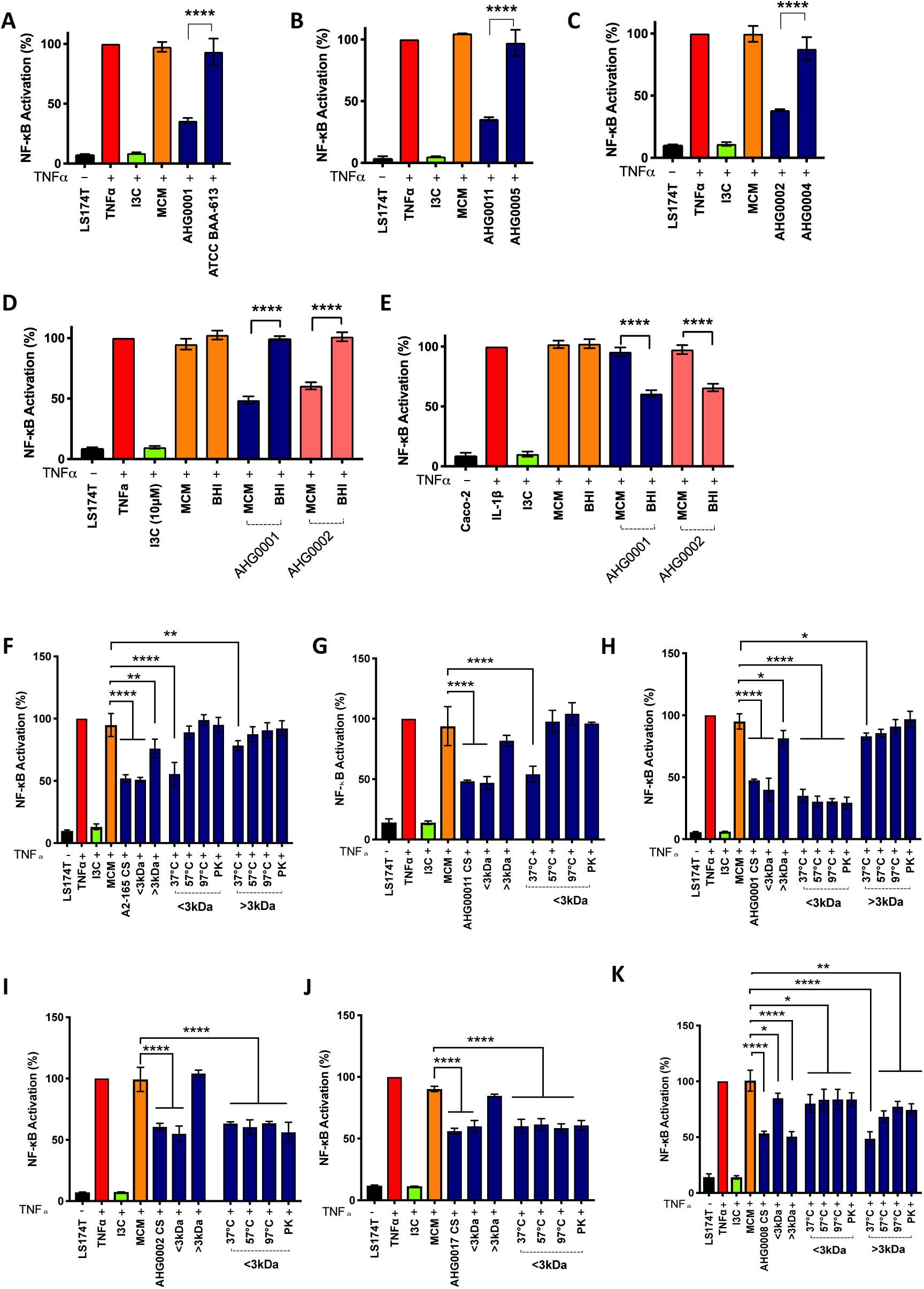
**A-C.** Characterisation of intraspecies variation in NF-κB suppressive capacity. The ability of *C. bolteae* AHG0001 and ATCC BAA-613 (Panel A), *C. citroniae* AHG0002 and AHG0004 (Panel B) and *C. aldenense* AHG0011 and AHG0005 (Panel C) to suppress NF-κB was analysed using the LS174T reporter cells. NF-κB activation was assessed 4 h after TNFα stimulation and the extent of suppression was assessed against sterile medium (mean (SD)). **D-E.** Characterisation of the effect of growth medium on the NF-κB suppressive capacity of *C. bolteae* AHG0001 and *C. citroniae* AHG0002 in LS174T (Panel D) and Caco-2 (Panel E) reporter cell lines. NF-κB activation was assessed 4 h after cytokine stimulation and the extent of suppression was assessed against sterile medium. **F-K.** Characterisation of the bioactive factors produced by *F. prausnitzii* A2-165 (Panel F), *C. aldenense* AHG0011 (Panel G), *C. bolteae* AHG0001 (Panel H), *C. citroniae* AHG0002 (Panel I), *E. limosum* AHG0017 (Panel J) and *Pseudoflavonifractor* sp. AHG0008 (Panel K). The cell free CS were untreated or subjected to size fractionation, heat and/or proteinase K treatments as appropriate. NF-κB activation was assessed 4 h after TNFα stimulation and the extent of suppression was assessed against sterile medium (mean (SD)). * *p*<0.05, ** *p*<0.01, *** *p*<0.001, **** *p*<0.0001 as determined by one-way ANOVA with Dunnett’s multiple comparison test.

We examined our collection of suppressive CS by a combination of size fractionation, proteinase K and heat treatments to determine their biochemical characteristics. Using this approach, we determined that the NF-κB suppressive activity for all strains except *Pseudoflavonifractor* sp. AHG0008 was predominantly associated with the <3 kDa fraction (Figure 2F-K, Supplementary Results). Gut bacteria produce a structurally diverse array of low molecular weight NF-κB suppressive bioactives^18, 20, 28^ and we focused on the <3 kDa fraction as we believed these bioactives would be more amenable to drug development. We concluded that these bioactives could be broadly separated into two classes based on heat and protease sensitivity (e.g. *F. prausnitzii* A2-165, *C. aldenense* AHG0011) or resilience (e.g. *C. bolteae* AHG0001, *C. citroniae* AHG0002, and *E. limosum* AHG0017), possibly inclusive of both peptides and/or thermal and hydrolytically stable small molecules, respectively (Figures 2F-J, Supplementary Results). We sequenced the strains producing <3 kDa bioactives to near completeness to identify candidate bioactive encoding biosynthetic gene clusters (BGCs) (Table 1). Phylogenetic analysis using the Genome Taxonomy Database (GTDB) confirmed the 16S rRNA based assignments (Supplementary Figure 2A). We also determined that the strains exhibited a high degree of genome synteny with their near relatives (Supplementary Figure 2B-E) and carried multiple BGCs (Table 1). None of the isolates encoded *F. prausnitzii* Mam-like orthologs which is consistent with its narrow phylogenetic distribution^18, 29^.

**Table 1.**

| **Genome designation** | **CheckM marker lineage (Order (GTDB Branch))** | **Number of contigs** | **Genome size (bp)** | **GC (%)** | **CheckM completeness (%)** | **CheckM contamination (%)** | **Number of BGC** | **NCBI accession number** |
| --- | --- | --- | --- | --- | --- | --- | --- | --- |
| *C. bolteae*<br>AHG0001 | Clostridiales (UID1342) | 96 | 5,985,600 | 49.4 | 98.76 | 0.56 | 19 | QYRW00000000 |
| *C. citroniae*<br>AHG0002 | Clostridiales (UID1226) | 149 | 6,630,634 | 48.8 | 99.37 | 0 | 25 | QYRX00000000 |
| *C. aldenense*<br>AHG0011 | Clostridiales (UID1226) | 263 | 6,734,822 | 49.5 | 99.37 | 0 | 24 | QYRY00000000 |
| *E. limosum*<br>AHG0017 | Clostridiales (UID1120) | 86 | 4,704,612 | 47.2 | 99.3 | 0.7 | 31 | QYRZ00000000 |

### CS suppress *ex vivo* IL-8 secretion

IBD is challenging to treat due to the variability of response to available medications. A proportion of this variability is related to underlying genetic susceptibilities which likely drive their evolved immunophenotype and host-microbiota relationship^30, 31^. To assess whether the suppressive CS could affect epithelial inflammatory responses in primary cells in the context of IBD associated genetic risk factors we assessed their ability to prevent IL-1β driven IL-8 production in healthy (n=6), CD (n=5) and UC (n=5) derived primary intestinal epithelial organoid cultures. Interestingly, despite removal from the inflammatory environment, there was significantly higher basal IL-8 production by organoids derived from CD patients compared to those from non-IBD controls and UC patients (Figure 3A). Following stimulation with IL-1β there was a significantly more IL-8 produced by organoids derived from UC but not CD when compared to healthy subjects (Figure 3B). As expected, IL-8 secretion was significantly inhibited by I3C, but not MCM in healthy, CD and UC subjects. Treatment with *F. prausnitzii* A2-165 CS resulted in significantly suppressed IL-8 secretion when compared to the MCM control in healthy and CD but not UC subjects (Figure 3C-E). Treatment with CS from *C. aldenense* AHG0011, *C. citroniae* AHG0002, *E. limosum* AHG0017, *C. bolteae* AHG0001 and *Pseudoflavonifractor sp.* AHG0008 significantly suppressed IL-8 secretion in healthy, CD and UC subjects, to an equivalent or greater degree than *F. prausnitzii* A2-165 (Figure 3C-E). There was a high degree of concordance in the degree of suppression between subjects within bacterial CS, in all subject groups, although some subject specific differences were noted (Supplementary Figure 3A). Critically, we did not observe any significant cytotoxic effects from the CS treatments (Supplementary Figure 3B-D).

**Figure 3.**
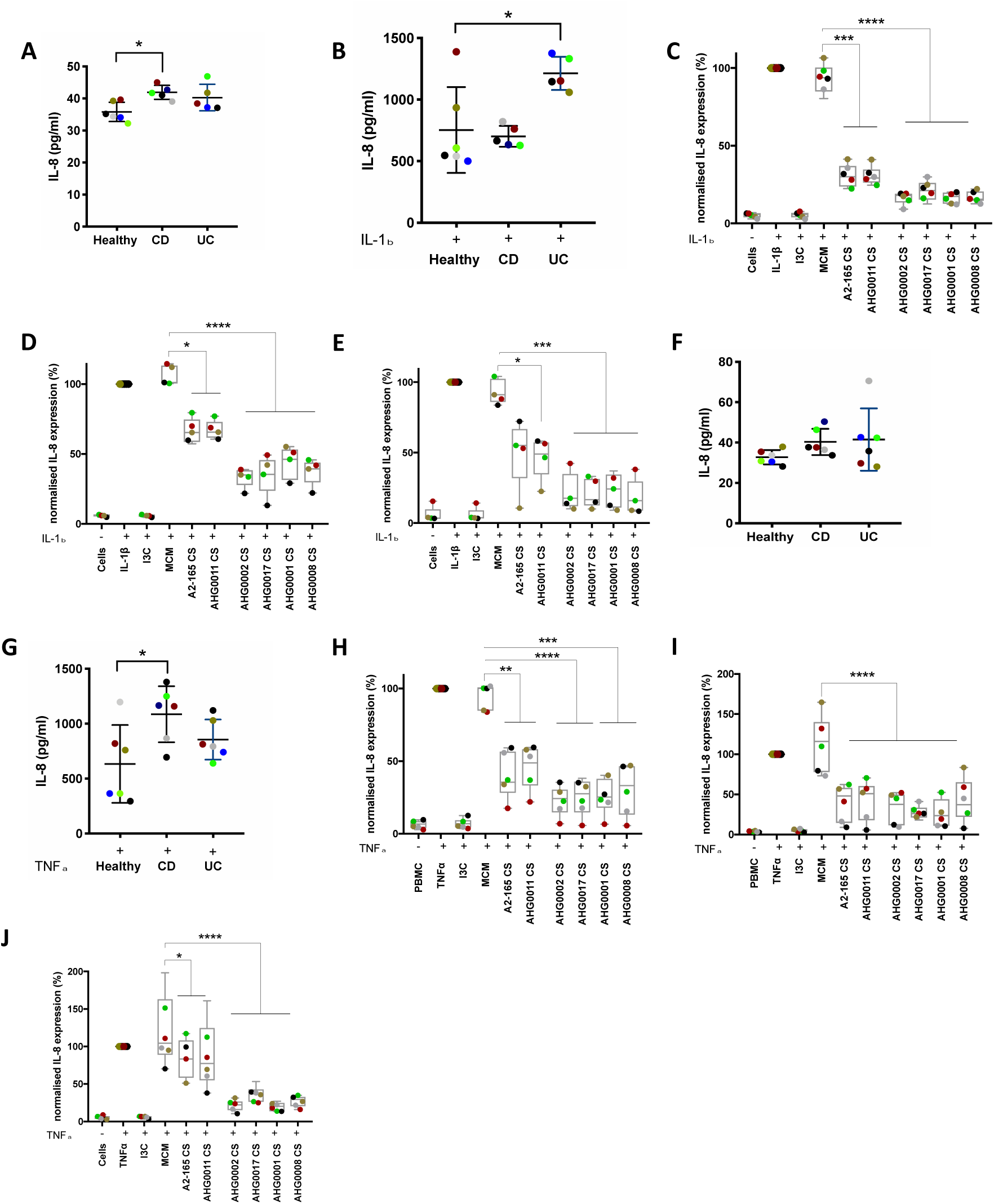
**A-B.** Analysis of basal IL-8 secretion by healthy, CD and UC derived gut epithelial organoids (Panel A) and following IL-1β stimulation (Panel B). IL-8 secretion was assessed 24 h after cytokine stimulation (mean (SD)). **C-E.** Analysis of the ability of *F. prausnitzii* A2-165, *C. aldenense* AHG0011, *C. citroniae* AHG0002, *E. limosum* AHG0017, *C. bolteae* AHG0001 and *Pseudoflavonifractor* sp. AHG0008 to suppress IL-8 secretion in healthy (Panel C), CD (Panel D) and UC (Panel E) subject derived gut epithelial organoids. IL-8 secretion was assessed 24 h after cytokine stimulation and compared against the sterile medium (mean (SD)). **F-G.** Analysis of basal IL-8 secretion by healthy, CD and UC derived PBMCs (Panel F) and following TNFα stimulation (Panel G). **H-J.** Analysis of the ability of *F. prausnitzii* A2-165, *C. aldenense* AHG0011, *C. citroniae* AHG0002, *E. limosum* AHG0017, *C. bolteae* AHG0001 and *Pseudoflavonifractor* sp. AHG0008 to suppress IL-8 secretion in healthy (Panel H), CD (Panel I) and UC (Panel J) subject derived PBMCs. IL-8 secretion was assessed 24 h after cytokine stimulation and compared against the sterile medium (mean (SD)). * *p*<0.05, ** *p*<0.01, *** *p*<0.001, **** *p*<0.0001 as determined by one-way ANOVA with Dunnett’s multiple comparison test.

In addition to effects on the epithelium, bioactives produced by gut bacteria may also be absorbed and have systemic effects on immune cells. Therefore, the suppressive effects of the CS on primary immune cells was examined using PBMCs collected from healthy, CD and UC (n=6 per group) subjects. While the basal concentrations of IL-8 released by PBMC from all three groups were not significantly different (Figure 3F), their stimulation with TNFα resulted in more IL-8 released from the PBMCs of the CD group in comparison to those prepared from the healthy or UC groups (Figure 3G). As expected, IL-8 secretion by PBMCs from healthy, CD and UC subjects was significantly inhibited by I3C and *F. prausnitzii* A2-165 CS (Figure 3H-J). Similarly, IL-8 secretion by PBMCs from healthy, CD and UC subjects was suppressed by treatment with CS from *C. aldenense* AHG0011, *C. citroniae* AHG0002, *E. limosum* AHG0017, *C. bolteae* AHG0001 and *Pseudoflavonifractor sp.* AHG0008; at least as effectively as *F. prausnitzii* A2-165 (Figure 3H-J). There was limited variation in the response to CS within the healthy, CD and UC subject groups although there were some subjects that showed varying levels of suppression with individual CS (Supplementary Figure 4A). Critically, we also did not observe any significant cytotoxic effects from the CS treatments on PBMCs (Supplementary Figure 4B-D). Collectively, these results show that the CS of these strains can suppress cytokine mediated inflammatory responses in the gut and immune compartments in both an IBD and non-IBD genetic background.

### Precision treatment of murine colitis

The NF-κB pathway is highly conserved in mammals and we next examined the ability of the CS to suppress IL-1β induced expression of the NF-κB regulated genes *Mip-2* and *Cxcl-10* in C57/Bl6 derived murine organoids. All the CS tested suppressed induction, suggesting that the bioactives likely act through conserved mammalian cell targets (Figure 4A-B). We also examined the ability of the CS to suppress expression of *Mip-2* and *Cxcl-10* in organoids derived from *Winnie* mice. *Winnie* mice carry a missense mutation in *Muc2* that results in protein misfolding, endoplasmic reticulum (ER) stress and defects in gut barrier function. These mice develop a spontaneous colitis characteristic of UC and are an excellent preclinical model for human treatments^32–34^. We found that the majority of CS significantly suppressed IL-1β induced expression of *Mip-2* and *Cxcl-10* on *Winnie* derived organoids. However, in contrast to the findings in wild-type organoids, CS from *C. aldenense* AHG0011 and *F. prausnitzii* A2-165 did not suppress *Mip-2* and *Cxcl-10* (Figure 4C-D). Furthermore, using *Winnie* derived gut epithelial organoids we determined *C. bolteae* AHG0001 but not *C. bolteae* ATCC BAA-613 CS suppressed induction of *Mip-2* and *Cxcl-10* expression, confirming the strain specific differences observed in the reporter cell lines (Figure 4E). Interestingly, we also determined that *C. bolteae* AHG0001 but not *C. bolteae* ATCC BAA-613 CS suppressed induction of the ER stress markers, *Grp78* and *sXbp1*, in *Winnie* organoids (Figure 4E).

**Figure 4.**
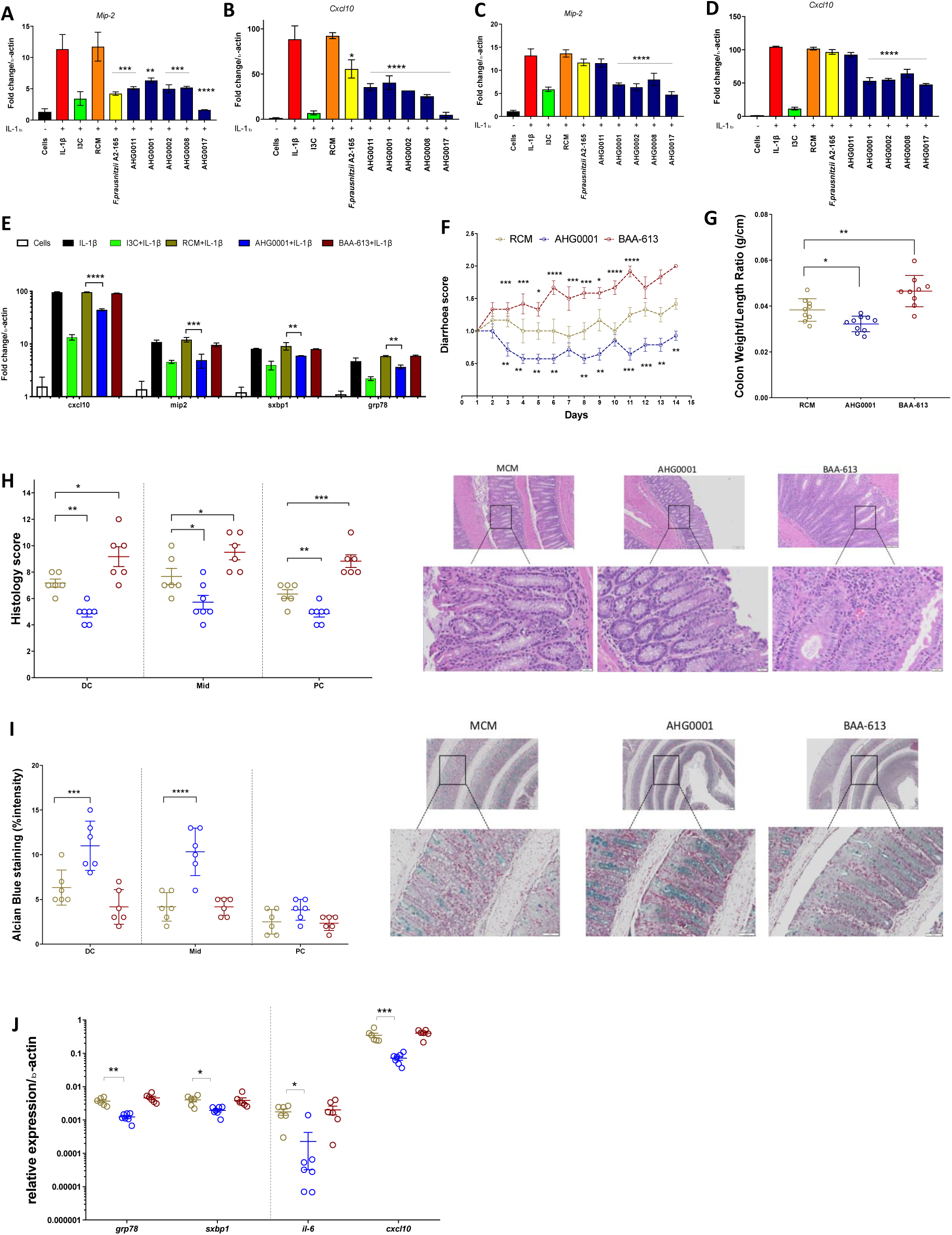
**A-D.** Effects of bioactives on pro-inflammatory gene expression using murine derived organoids from C57/BL6 (Panels A-B) and *Winnie* (Panels C-D) mice. Murine derived organoids were treated with CS for 30mins and then stimulated as appropriate with mIL-1β for 6 hours. **E.** *Winnie* organoid based qRT-PCR quantification of *cxcl10*, *mip-2*, *sxbp1* and *grp78* expression following treatment with *C. bolteae* AHG0001 or *C. bolteae* BAA-613. **F.** Effect of daily administration of MCM, *C. bolteae* AHG0001 CS or *C. bolteae* BAA-613 CS on diarrhoea score. **G.** Changes in colon weight/length ratio following treatment with MCM, *C. bolteae* AHG0001 CS or *C. bolteae* BAA-613 CS. **H.** Blinded histology scores following treatment with MCM, *C. bolteae* AHG0001 CS or *C. bolteae* BAA-613 CS. **I.** Alcian blue quantification of mucin production in *Winnie* derived colon sections with representative images from distal colon. **J.** Relative gene expression of ER stress markers (*grp78 and sxbp1*) and pro-inflammatory (*il-6*, *cxcl10*) genes in colonic tissue sections as analysed by qRT-PCR. ns not significant, * *p*<0.05, ** *p*<0.01, *** *p*<0.001, **** *p*<0.0001. The significance for diarrhoea was determined by comparison with MCM using one-way ANOVA with Dunnett’s multiple comparison test. Sidak’s multiple comparison tests were used for Figures E-J.

We hypothesised that functional capacity rather than phylogeny would be the principle determinant of therapeutic efficacy and that primary organoid cultures could be used to predict *in vivo* host responses to select CS in a precision medicine-based manner. To test this, CS prepared from *C. bolteae* AHG0001 and ATCC BAA-613 were administered intrarectally for 14 days to 6-week old *Winnie* mice with established colitis, as demonstrated by the elevated diarrhoea scores at the start of the experiment (Figure 4F). *C. bolteae* AHG0001 CS significantly reduced diarrhoea scores over the course of the experiment compared to MCM and *C. bolteae* ATCC BAA-613 CS treated animals (Figure 4F). Furthermore, CS from *C. bolteae* AHG0001 significantly reduced colonic inflammation as determined by a decreased colon weight to length ratio (Figure 4G), histology scores (Figure 4H, Supplementary Figure 5A-C) and immune cell infiltration (Supplementary Figure 5E). Moreover, *Winnie* mice treated with *C. bolteae* AHG0001 demonstrated increased mucin production and goblet cell restitution in the distal and mid colon as determined by Alcian blue staining (Figure 4I, Supplementary Figure 5D); indicative of reduced endoplasmic reticulum (ER) stress and histologic healing. Consistent with reduced colitis, there was a significant reduction in colonic expression of the inflammatory genes *Il-6* and *Cxcl-10* and the ER stress markers *spliced-Xbp1* and *Grp78* in the colon (Figure 4J). Together, these results showed the feasibility of applying a precision medicine approach using *ex vivo* organoid cultures to accurately predict treatment response in colitis.

Finally, we hypothesised that intraspecies variations in NF-κB suppressive capacity, together with the influence of culture media on bioactive production, could facilitate identification of candidate bioactive encoding BGCs and/or bioactive molecules produced by *C. bolteae* using comparative genomics or metabolomics. Comparative genomic analyses revealed that *C. bolteae* AHG0001 carries 19 predicted BGCs of which 14 are either highly or partially conserved in *C. bolteae* ATCC BAA-613 (Figure 5A, Supplementary Figure 6A). However, as the biosynthesis of bioactives by gut bacteria may be principally driven through modest modifications of common primary metabolites^35, 36^ we considered it likely that other BGCs would be overlooked by *in silico* screens. We consequently applied a process of bioassay guided solvent extractions and filtrations, followed by ultra-high-performance liquid chromatography quadrupole time-of-flight mass spectrometric analysis (UPLC-QTOF), and comparative metabolomics, to identify the bioactive(s) (Figure 5B, Supplementary Information). These analyses successfully identified a cluster of six structurally related small molecules (Figure 5C, 5Ca, i-vi) that were uniquely present in the NF-κB suppressive ethyl acetate (EtOAc) extract of *C. bolteae* AHG0001, but were absent in comparable extracts of *C. bolteae* ATCC BAA-613, and, following semi-preparative HPLC fractionation of the *C. bolteae* AHG0001 EtOAc extract, were uniquely localised in the NF-κB suppressive fractions (Supplementary Figure 6B-C). The identification of a novel secreted molecule confirmed the benefit of using an integrated approach combining bacterial isolation, functional screens and comparative metabolomics to expedite bioactive discovery.

**Figure 5.**
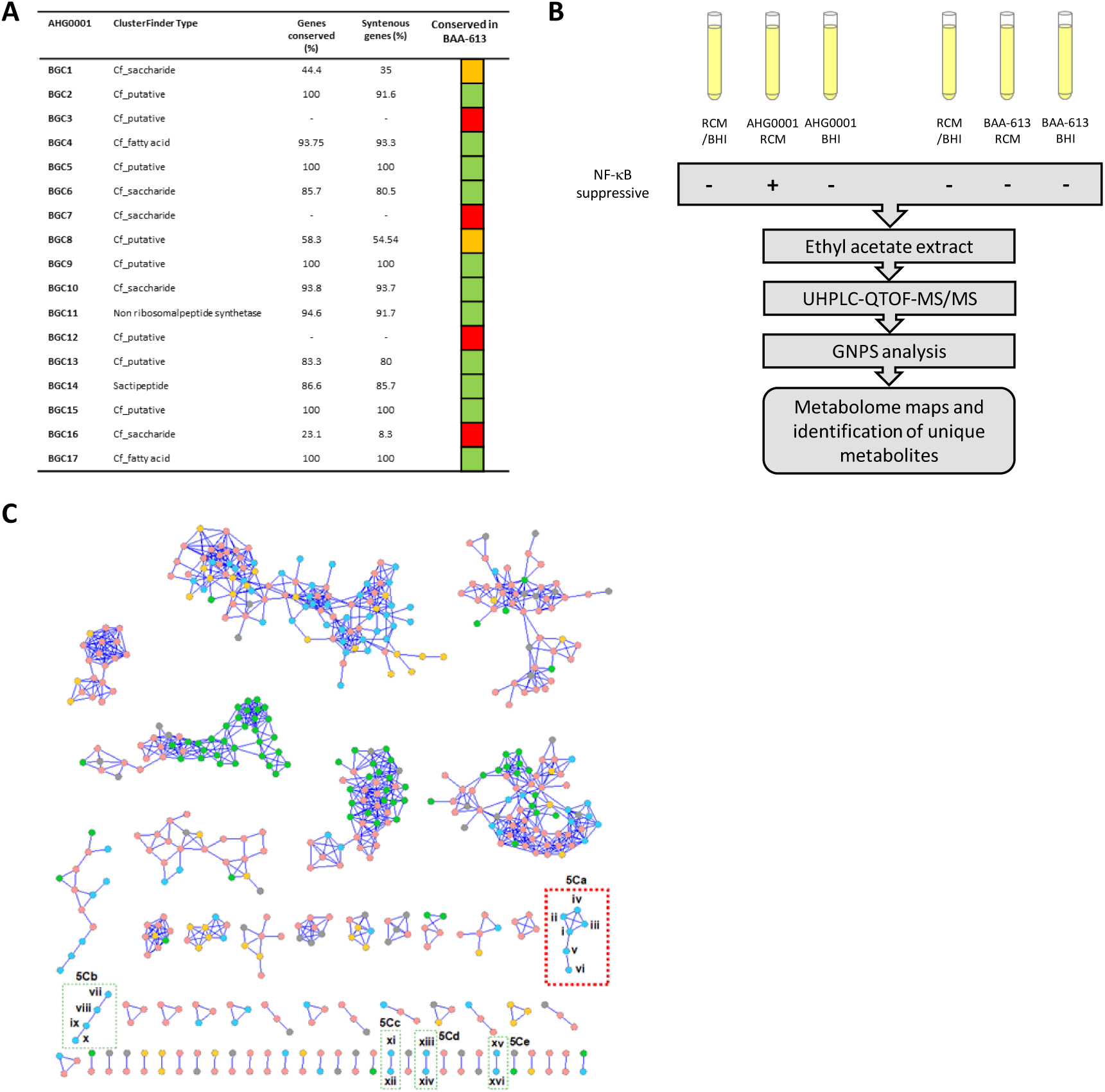
**A.** Determination of the extent of *C. bolteae* AHG0001 BGC conservation in *C. bolteae* ATCC BAA-613. The extent of protein (Genes conserved) and syntenic gene pair (Syntenous pairs) conservation was assessed. *C. bolteae* AHG0001 BGC were classed as being conserved (green), partially conserved (orange) or not conserved (red). **B.** An overview of the experimental approach used to identify bioactives associated with the NF κB suppressive activity of *C. bolteae* AHG0001. The presence of the bioactive in the various extractions and filtrates was determined using the LS174T reporter cell assay. **C.** Molecular networking for EtOAc extracts of *C. bolteae* AHG0001 cultured in MCM (blue nodes) and BHI media (green nodes); yellow and grey nodes represent compounds from MCM and BHI media only respectively; pink nodes represent compounds common to MCM and BHI extracts or MCM and BHI media. Boxes 5Ca, 5Cb, 5Cc, 5Cd and 5Ce highlight structurally related small molecules clusters (i-vi, vii-x, xi-xii, xiii-xiv and xv-xvi respectively), that are unique to the NF-κB suppressive EtOAc extracts of *C. bolteae* AHG0001 cultured in MCM media. Only the molecules within the 5Ca cluster (red dashed box) are present in semi-preparative HPLC fractions that exhibit NF-κB suppressive activity.

## Discussion

Firmicutes affiliated bacteria are amongst the most abundant gut microbes and these taxa are widely recognised to possess immunomodulatory capacities^15, 37, 38^. However, they are poorly represented in culture collections and their ability to modulate immune responses remain largely undefined. In this study, we identified five gut bacterial strains affiliated with *Clostridium* clusters IV, XIVa and XV that are comparable to the well-characterised *F. prausnitzii* A2-165 strain in their ability to suppress NF-κB activation. The NF-κB suppressive bioactivities were characterised by significant biochemical and intraspecies variations suggesting there may be extensive functional redundancy and NF-κB suppressive capacity may be more prevalent than previously appreciated. This is consistent with Geva-Zatorsky *et al*.,^2^ who determined that as few as 53 isolates were associated with over 24,000 immune phenotypes that include functionalities relevant to IBD (e.g. Treg induction). Modulating host immune responses may support the ability of gut bacteria to colonise and persist in the gut environment and the ability of the microbiota to act as an extrinsic regulator of host immunity may underpin immune homeostasis and contribute to disease risk in genetically susceptible individuals.

IBD is characterised by a dysregulated immune response with select genetic susceptibilities affecting therapeutic responsiveness^30, 31^. In order to develop improved precision treatments for IBD we therefore used gut epithelial organoids and immune cells to identify bacteria capable of supressing cytokine mediated inflammatory responses. The heat and proteinase K resilient bioactives showed strong suppression of IL-8 secretion in organoids and immune cells from healthy, CD and UC subjects. Interestingly, the putative peptide bioactives produced by *F. prausnitzii* A2-165 and *C. aldenense* AHG0011 were notably less suppressive in UC derived organoids and PBMCs, and CD organoids, when compared to organoids derived from healthy controls; this may be reflective of the increased endogenous protease activity in IBD^39^. Our *in vitro* and *ex vivo* data also suggested that functional capacity rather than phylogeny may be the key determinant of biologic effects. To explore this hypothesis, we capitalised on the *C. bolteae* intraspecies differences and demonstrated that a precision medicine approach could be applied to alleviate established colitis in *Winnie* mice. Notably, treatment with *C. bolteae* AHG0001 CS was associated with a rapid onset of action with improvement in diarrhoea, alleviation of inflammation and ER stress, as well as restoration of goblet cell numbers and mucin production. Mucosal and histologic healing are amongst amongst the best predictors of long-term outcomes in IBD and taken together our data suggests a precision medicine approach could be applied to microbiome based IBD treatment.

The NF-κB suppressive strains carry multiple BGCs, many of whose products remain cryptic, underlining the inherent challenges in applying genomic based approaches to map genotype with phenotype. In addition, the biosynthesis of bioactives by gut bacteria may be principally driven through modest modifications of common primary metabolites that are underpinned by small BGCs^35, 40^. As the medium dependent effects on NF-κB suppression may affect the therapeutic efficacy of live biotherapeutics for IBD, we therefore used a bioassay-guided fractionation and a comparative metabolomic approach to identify a novel low molecular weight non-polar molecule that was associated with the NF-κB suppressive activity of *C. bolteae* AHG0001. Critically, this molecule was not associated with medium components suggesting the suppressive activity is unlikely to be due to the biotransformation of medium components^41^. Consistent with other microbial bioactives, the *C. bolteae* AHG0001 bioactive acts independently of the bacterial cell and suppresses the inflammatory response in animals.

In summary, our IBD guided approach provides new opportunities to rationally bioprospect the gut microbiota for precision live biotherapeutic strains and/or bioactives that could be used to expedite the development of safer and more efficacious therapeutics.

## Materials & Methods

### Bacterial strains, culture conditions and analyses

Firmicutes affiliated bacteria were cultured in anoxic MCM or BHI medium (Supplementary Information). *E. coli* Nissle 1917 was cultured using LB medium. *L. casei* Shirota was isolated from a Yakult Original probiotic drink and cultured using anoxic de Man Rogosa Sharpe (MRS) or BHI medium. *L. rhamnosus* GG and *B. animalis* subsp. *lactis* BB-12 were isolated from a probiotic capsule and cultured using anoxic MRS or BHI medium.

### Bacterial comparative analyses

The phylogeny of the Firmicutes isolates was inferred using the *rrs* gene sequences as described in the Supplementary Information. High molecular weight DNA was prepared and sequenced using the Illumina NextSeq 500 system (2 x 150bp High Output kit) with v2 chemistry as previously described^20^. The *C. bolteae* AHG0001, *C. citroniae* AHG0002, *C. aldenense* AHG0011 and *E. limosum* AHG0017 sequence data were assembled, assessed for contamination and completeness and ordered as described in the Supplementary Information. Genome based phylogeny was determined using the Genome Taxonomy Database (GTDB) as previously described^20^. Candidate BGC were identified and characterised as described in the Supplementary Information. The Whole Genome Shotgun projects for *C. bolteae* AHG0001, *C. citroniae* AHG0002, *C. aldenense* AHG0011 and *E. limosum* AHG0017 were deposited respectively at DDBJ/EMBL/GenBank under the accessions QYRW00000000, QYRX00000000, QYRY00000000 and QYRZ00000000. The version described in this paper are the first versions, [XXXX]01000000.

### Measurement of immunomodulatory activities

The immunomodulatory potential of the individual strains was examined following growth in MCM, BHI or MRS. Briefly, for the first pass screen an individual colony was used to inoculate medium and the culture was grown for up to 96 hours. For the confirmatory screens, the NF-κB suppressive capacity of biological replicate cultures produced from select strains was assessed. Briefly, two independent broth cultures were established from individual colonies of each strain (n=2 independent biological replicates per strain) and following growth as described above, each individual culture was used to inoculate 3 tubes of broth (n=6, consisting of n=2 independent biological replicates per strain with n=3 technical replicate for each biological replicate). The cultures were grown until early stationary phase and then 1.5 ml of each culture was centrifuged at 25,000 x g for 3 minutes. Then, 1 ml of the cell-free supernatant fraction was collected and stored at ≤30°C as a single-use aliquot. The NF-κB suppressive capacity of the CS was assessed using the LS174T-NF-kB*luc* or Caco-2-NF-kB*luc* reporter cell assays adapted for high-throughput screening (Supplementary Information). The effects of sodium salt SCFA on NF-κB activation were assessed by treating the cell lines for 30 min and then determining their ability to suppress cytokine mediated NF-κB activation as described above. The cytotoxicity of the supernatants was assessed using the CytoTox96® Non-Radioactive Cytotoxicity Assay according to the manufacturer’s instructions (Promega).

### Organoid culturing and immunomodulatory assays

All patient samples were collected in accordance with the recommendations of the Mater Health Services Human Research Ethics Committee (HREC 2016001782 & HREC/14/MHS/125) for the Mater Inflammatory Bowel Disease Biobank. Colonic biopsies (6 x 3mm pinch biopsies) were collected from healthy (n=6), CD (n=5) and UC (n=5) patients (Supplementary Table 1). The colonic biopsies were processed and cultured as previously described (Supplementary Information). To assess the ability of the CS to suppress IL-8 secretion the organoids were seeded in a 48 well plate and grown for 48 hours. Then, organoids were treated with 10% v/v of select CS in 50% L-WRN conditioned medium and subsequently stimulated with rhIL-1β (50 ng.ml^-1^) for 24 hours before quantifying IL-8 in the supernatant. Cytotoxicity was assessed using the CytoTox 96® Non-Radioactive Cytotoxicity Assay. For the animal experiments, colonic tissues from C57BL/6 and *Winnie* (n=2) mice were segmented and the crypts were isolated and cultured according to previously established protocols (Supplementary Information). For the treatments, the organoids were first seeded in a 24 well plate and grown for 48 hours. The organoids were then pre-treated with 10% v/v of select CS for 30 mins and then stimulated with 50 ng/ml mIL-1β for 6 hours. The cells were lysed and used for mRNA expression.

### Peripheral Blood Mononuclear Cell (PBMC) isolation and immunomodulatory assays

Human peripheral blood was obtained for 6 healthy, CD and UC patients from the Mater Inflammatory Bowel Disease biobank. PBMCs were isolated by Ficoll gradient density centrifugation (Supplementary Information). For the treatments, 500,000 cells per well were plated on a 96-well plate and treated with 10% v/v of CS in RPMI medium for 30 minutes, followed by stimulation with rhTNFα (50 ng/ml). IL-8 secretion and cytotoxicity was assessed as previously described.

### RNA extraction, cDNA synthesis and gene expression

Total RNA was prepared as previously described except that LS174T and Caco-2 cells were used. The expression of *il6*, *il8* and *cxcl-10* was assessed as previously described^20, 21^. RNAeasy mini kits (QIAGEN) were used to extract RNA from the mouse organoid cultures according to the manufacturer’s instructions. RNA was reverse transcribed to cDNA using iScript cDNA synthesis kit (Bio-Rad Laboratories) and the protocol provided by the manufacturer. The Ct values for each gene were normalised to untreated controls and further normalised to housekeeping gene (mouse β-actin) and presented as fold change. The mouse primers used are summarised in Supplementary Table 2).

### Quantitative cytokine expression assays

To quantify IL-8 secretion, cell free supernatant was collected after 24 hours and IL-8 was quantified by ELISA according to manufacturer’s instructions (BioLegend).

### Animal experiments

All animal experiments were approved by the University of Queensland Animal Ethics Committee. *Winnie* mice were bred in-house in a pathogen-free animal facility. Male and female mice were intrarectally gavaged with 50 µl of CS from *C. bolteae* AHG0001 and ATCC BAA-613 for 14 days. MCM medium processed in the same manner as the CS was used as the vehicle control. Disease activity was assessed as described in the Supplementary Information.

### GNPS Analyses

UHPLC-QTOF (Agilent Technologies 6545 Q-TOF LC/MS) data was acquired by subjecting aliquots of EtOAc extracts obtained from (a) cultures of *C. bolteae* AHG0001 in either MCM or BHI media, (b) cultures of *C. bolteae* BAA-613, in either MCM or BHI media, and; (c) un-inoculated MCM and BHI media (1 μL). UHPLC conditions were as described in the Supplementary Information. The acquired MS/MS data was converted from Agilent MassHunter data file (.d) to the mzXML file format using the software MS-Convert^42^. Molecular networks were generated using the online Global Natural Products Social molecular networking web-platform (GNPS) (gnps.ucsd.edu). MS-Cluster with a precursor ion mass tolerance of 0.02 Da and a MS/MS fragment ion tolerance of 0.02Da were selected to create consensus spectra^43^. A minimum cluster size of 1, cosine score 0.7, and minimum number of fragments of 6, were selected for molecular networking. The spectral networks were imported into Cytoscape 3.5.1^44^ and visualized using force directed layout where nodes represented parent masses.

### Analytical fractionation of NF-κB suppressive extract

An EtOAc extract (3mg) of *C. bolteae* AHG0001 cultivated in MCM medium was subjected to analytical HPLC (Supplementary Information) to yield 17 fractions. Each fraction was dried *in vacuo* then resuspended in MeOH (50 μL). NF-κB suppressive fractions were combined and an aliquot (1 μL) was subjected to UHPLC-QTOF analysis, with single ion extraction (SIE) same quadrupole time-of-flight spectrometer and UHPLC conditions described above. Single ion extraction (*m/z* molecular ion) chromatograms for molecules exclusively present in *C. bolteae* AHG0001 (i.e. i-xvi in GNPS analysis Figure 5C), using Agilent MassHunter Qualitative Analysis software, confirmed that only i-vi were present in the NF-κB suppressive fraction.

### Statistical analyses

The NF-κB suppressive effects of the suppressive strains was assessed using biological duplicates with each duplicate comprising of three technical replicates. Significance was determined using a one-way ANOVA with correction for multiple comparisons with a Dunnett test. Differences were considered significant at *p*≤0.05. A heat map of the first pass screen data was produced using GraphPad Prism (version 7.0) and the Heatmap tool at the HIV sequence database (https://www.hiv.lanl.gov/content/sequence/HEATMAP/heatmap.html). The animal experiments were performed twice independently, and the data combined for analysis. The D’Agostino-Pearson omnibus test was used to verify the normal distribution of all data. Significance was determined using t-tests, one-way ANOVA with multiple comparisons (Sidak), two-way ANOVA corrected for multiple comparisons with a Dunnett test using GraphPad Prism (version 7.0). Differences were considered significant at *p* ≤0.05.

## Acknowledgements

This research was supported via funds provided by the University of Queensland (UQ) Faculty of Medicine (MM, JB and PÓC) and Diamantina Institute (MM). We gratefully acknowledge the support provided by the UQ Research Training Program and Mater Frank Clair Scholarship (RG), UQ Institute for Molecular Bioscience (KS and RKC) and UQ Reginald Ferguson Fellowship in Gastroenterology (PÓC). The Translational Research Institute is supported by a grant from the Australian Government.

## Author Contributions

PÓC and JB conceived the study and developed it with MMcG and MM; PÓC and RG prepared samples for analysis and performed the immunomodulatory characterizations; RG and JB performed the organoid, PBMC and animal experiments; ECH and PÓC performed the genomic analyses; KS and RJC performed the metabolomics and molecule analyses; RG, ECH, KS, MMcG, MM, RJC JB and PÓC analysed the data, and; PÓC wrote the manuscript with RG, ECH, KS, MMcG, MM, RJC and JB.

## Competing Interest

The authors declare no competing interest.

## Supplementary Results

### Bioactive identification

Based on the strain and medium effects on *C. bolteae* NF-κB suppressive activity we applied a comparative metabolomics approach to identify the bioactive. During purification we noted that direct filtering of CS through a 0.42 µm nylon filter resulted in a loss of NF-κB suppressive activity, suggestive of either low water solubility and/or a non-polar bioactive(s). By contrast, an ethyl acetate (EtOAc) extract derived from the same CS was readily filtered, with the filtrate retaining NF-κB suppressive activity. As a next step in the chemical characterisation, EtOAc extracts were prepared from *C. bolteae* AHG0001 and *C. bolteae* BAA-613 following growth in MCM and BHI along with EtOAc extracts from both un-inoculated MCM and BHI media. Each extract was individually subjected to ultra-high-performance liquid chromatography quadrupole time-of-flight mass spectrometric analysis with MS/MS monitoring (UPLC-QTOF-MS/MS), followed by global natural products social molecular networking (GNPS) analysis (Figure 5B), to generate a metabolome map with media controls. These analyses revealed multiple clusters of metabolites that appeared unique to the NF-κB suppressive MCM CS extract however only a single cluster of 6 novel and previously unreported metabolites (i-vi) was not co-clustered with compounds present in culture media, or the non-suppressive BHI CS extract. To confirm whether metabolites i-vi were the target bioactives, a portion of the NF-κB suppressive EtOAc extract was subjected to fractionation through a reversed-phase analytical HPLC column, with timed collection of 14 fractions. Significantly, NF-κB suppressive activity was localised in the non-polar fractions #13 and #14, which UPLC-QTOF analysis using single ion extraction (SIE) monitoring confirmed to co-localise with metabolites i-vi. By contrast, UPLC-QTOF-SIE analysis of other compounds present in media-associated clusters (Figure 5B), revealed they did not uniquely co-localise into the active fractions.

## Supplementary Methods & Materials

### Bacterial strains, culture conditions and analyses

Anaerobic Firmicutes affiliated bacteria were cultured in anoxic MCM (Lab-Lemco 10 g.L^-1^, Peptone P 10 g.L^-1^, Yeast extract 3 g.L^-1^, Glucose 5 g.L^-1^, Starch 2 g.L^-1^, Sodium chloride 5 g.L^-1^, Sodium bicarbonate 15 g.L^-1^, Resazurin 1 mg.L^-1^, Cysteine-HCl 1 g.L^-1^) or BHI supplemented with salt solutions 2 and 3^1^. *F. prausnitzii* A2-165 was grown as previously described^2^. A Coy vinyl anaerobic chamber with an anoxic atmosphere (85% N2:10% CO2:5% H2) was used to process the anaerobic Firmicutes cultures. Bacterial cultures were incubated at 37°C for up to 48 hours. Bacterial growth was measured by spectrophotometry (OD600nm) using a SPECTRONIC 20D+ Spectrophotometer (ThermoFisher, Sydney).

### Bacterial comparative analyses

Phylogenetic trees were constructed by aligning the 16S rRNA gene sequences using the SILVA database^3^ and the alignment was then imported into MEGAX^4^. The alignment was refined, and a maximum-likelihood phylogenetic tree constructed displaying the isolate and select reference sequences. The stability of the maximum-likelihood tree was evaluated by 1000 bootstrap replications and Kimura 2-parameter modelling. Where necessary, select isolates were subject to whole cell protein profiling to determine intraspecies variations^5, 6^. High molecular weight DNA was prepared as previously described^7^. The SPAdes assembler v 3.11.0 was used to quality check, filter and then *de novo* assemble the sequence data^8^. CheckM^9^ was used to evaluate the genome sequencing quality by estimating the completeness and contamination based on the phylogenetic assignment of a broad set of marker genes. The *C. bolteae* AHG0001, *C. citroniae* AHG0002, *C. aldenense* AHG0011 and *E. limosum* AHG0017 contigs were ordered using Mauve^10^ with the *C. bolteae* ATCC BAA-613, *C. citroniae* WAL-17108, Clostridiales bacterium 1_7_47FAA and *E. limosum* ATCC 8486 genome sequences respectively as references. Genome based phylogeny was determined using GTDB^11^ as previously described^2^. Candidate BGC were identified using the antiSMASH webserver^12^ with the ClusterFinder Detection Strictness settings set to “loose” and the Extra Features turned on. Similar candidate BGC were identified in select genomes or the Genbank Database using MultiGeneBlast^13^ in homology search mode. BGCs were considered highly conserved if (i) ≥80% of the genes in an *C. bolteae* AHG0001 BGC were conserved in *C. bolteae* ATCC BAA-613, with genes defined as being conserved if the query exhibited ≥80% sequence identity over ≥80% of the query length, and; (ii) ≥ 70% of the potential syntenic genes in a *C. bolteae* AHG0001 BGC were conserved in a *C. bolteae* ATCC BAA-613 BGC (calculated as ((MultiGeneBlast Total score – No. of Blast hits)/0.5)/(No. of syntenic genes in *C. bolteae* AHG0001 BGC)). BGC were considered partially conserved if ≥40% of both the genes and potential syntenic genes were conserved.

### Measurement of immunomodulatory activities

The LS174T-NF-kB*luc* or Caco-2-NF-kB*luc* reporter cell lines were adapted for high-throughput screening using the criterion defined by Zhang *et al*.,^14^ where a Z-factor ≥0.5 represents an excellent assay, thereby providing a sensitive and specific approach to assess the NF-κB suppressive capacity of the isolates. The Z-factor for each assay was determined and only assays achieving a Z-factor ≥0.5 were processed for further analysis. The high-throughput assays were performed in 96-well microtiter plates as previously described^2^ except that the LS174T reporter cells were stimulated with 50 ng.ml^-1^ TNFα and the Caco-2 cell lines were treated with 7.5% v/v CS in complete DMEM medium. NF-κB driven luciferase expression was assessed using the Pierce^TM^ Firefly Luc One-Step Glow Assay Kit (ThermoFisher Scientific) according to the manufacturer’s instructions. The NF-κB suppressive isolates were scored and ranked on their Z-score^15, 16^.

### Organoid culturing and immunomodulatory assays

The colonic biopsies were processed and cultured as previously described^17^. Briefly, the biopsies were washed with PBS and digested with collagenase type I (2 mg.ml^-1^) supplemented with gentamicin (50 µg.ml^-1^) for 15-20 minutes at 37°C. The isolated crypts were washed with DMEM/F12 medium and centrifuged at 50 x g for 5 mins at 4°C. The pellets were then suspended in Basement Membrane Extract (BME, Invitrogen) in a 1:1 ratio. Then, 20µl of the mixture was plated in a 24 well tissue culture plate and cultured in 50% L-WRN conditioned medium. The crypts were expanded by serial culture until sufficient numbers were obtained for experimentation. To assess the ability of the CS to suppress IL-8 secretion the organoids were seeded in a 48 well plate and grown for 48 hours. Then, organoids were treated with 10% v/v of select CS in 50% L-WRN conditioned medium for 30 min and subsequently stimulated with rhIL-1β (50 ng.ml^-1^) for 24 hours before quantifying IL-8 in the supernatant. Cytotoxicity was assessed using the CytoTox 96® Non-Radioactive Cytotoxicity Assay. Colonic tissues from C57BL/6 and Winnie mice (n=2) were segmented and the crypts were isolated and cultured. Briefly, the tissues were segmented and washed with PBS, followed by EDTA (8mM) digestion for 1 hour at 4°C and further digested with collagenase type I (2 mg.ml^-1^) (Thermo Fisher Scientific) supplemented with gentamicin (50 µg.ml^-1^) for 15-20 minutes at 37°C. The isolated crypts were washed with complete F12 medium (Identical to complete media except DMEM/F12 was used instead of DMEM) and centrifuged at 50 x g for 5 mins at 4°C. The pellets were then suspended in BME in a 1:1 ratio. Then, 20µl of the mixture was plated in a 24 well tissue culture plate and cultured in 50% L-WRN conditioned medium. The crypts were expanded by serial culture until sufficient numbers were obtained for experimentation.

### Peripheral Blood Mononuclear Cell (PBMC) isolation and immunomodulatory assays

PBMCs were isolated by Ficoll gradient density centrifugation. Briefly, 20 ml of freshly drawn blood was diluted in phosphate buffered saline (1:2) and well mixed. The diluted blood was then carefully layered over Ficoll paque. The tubes were centrifuged without brakes at 400g for 20 minutes at 20°C. The interphase containing mononuclear cells were transferred into a new tube and washed twice in PBS. Prepared cells were stored in liquid nitrogen until required.

### Animal experiments

Disease activity was assessed using established protocols. Briefly, the body weights of the mice as well as diarrhoea and rectal bleeding were monitored and recorded daily. Diarrhoea scoring was interpreted as follows: 0 = no diarrhoea, solid stool; 0.5 = very mild diarrhoea, moist but formed stool; 1 = mild diarrhoea, formed but easily bisected by pressure applied with pipette tips; 1.5 = diarrhoea, no fully formed stools, and; 2 = severe, watery diarrhoea with minimal solid present. For histology scoring, the whole colon was rolled, fixed in 10% neutral buffered formalin, and paraffin embedded and sectioned and stained with Haematoxylin and Eosin (H&E) and Alcian blue. Blind assessment of histologic inflammation (increased leukocyte infiltration, neutrophil counts, depletion of goblet cells, crypt abscesses, aberrant crypt architecture, increased crypt length, and epithelial cell damage and ulceration) for proximal, mid and distal colon was performed as previously described. To quantify *in vivo* gene expression, the distal colon was snap frozen and homogenised in TRIzol. RNA was extracted using the Bioline RNA extraction kit according to manufacturer’s instructions. RNA concentration was measured using a Nanodrop 1000 spectrophotometer, followed by cDNA synthesis using 1 µg of RNA and the iScript cDNA synthesis kit (BioRad). The expression of genes of interest (Supplementary Table 2) were analysed using quantitative real time PCR (qRT-PCR) as previously described^24, 25^. C_t_ values were generated, and relative quantitation was determined by the ΔC_t_ method.

### GNPS Analyses

UHPLC conditions involved 0.5 mL.min^-1^ gradient elution from 10% CH3CN/H2O to 100% CH3CN over a period of 4.5 min, with constant 0.1% formic acid, through an Agilent SB-C_8_ 1.7 μm, 2.1 × 150 mm column (Agilent Technologies Inc., Mulgrave, VIC, Australia). The source parameters were: electrospray positive ionisation; mass range of m/z 50-1700; scan rate 10 × per sec; MS/MS scan rate 3 × per sec; fixed collision energy 40 eV; source gas temperature 325° C; gas flow 10 L.min^-1^; and nebulizer 20 psig. The scan source parameters were: VCap 4000; fragmentor 100; skimmer 45; and octopole RF Peak 750.

### Analytical fractionation of NF-κB suppressive extract

An EtOAc extract (3 mg) of *C. bolteae* AHG0001 cultivated on MCM medium was subjected to analytical HPLC (Agilent Zorbax SB-C8, 5 μm, 4.6 mm×150 mm column, gradient elution at 1 mL.min^-1^ from 10% MeCN/ H_2_O to 100% MeCN over 15 min followed by 2 min wash with 100% MeCN, without TFA modifier) to yield 17 fractions. Only fractions 14-17 demonstrated an ability to suppress NF-κB activity.

---

**Supplementary Figure 1.**
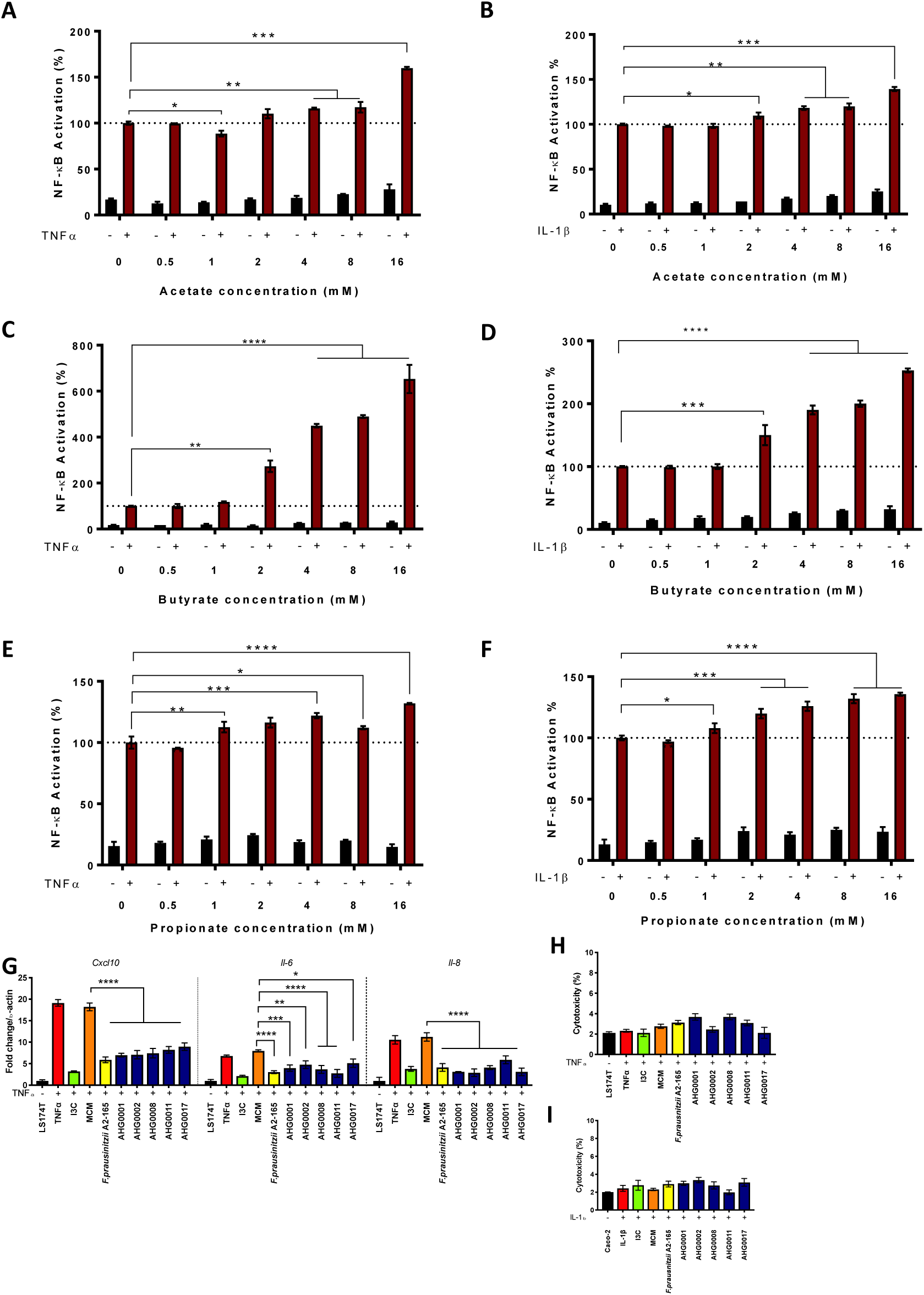
**A-F.** Assessment of the ability of acetate (Panels A-B), butyrate (Panels C-D) or propionate (Panels E-F) to suppress NF-κB activation in unstimulated or cytokine stimulated LS174T and Caco-2 reporter cells. NF-κB activation was assessed 6h after TNFα stimulation and the extent of suppression was assessed against sterile medium (mean (SD)). **G.** LS174T based qRT-PCR confirmatory assay of the hits identified from the first pass screen (mean (SD)). *F. prausnitzii* A2-165 and the validated hits suppress IL-1β induced *cxcl10*, *il6* and *il8* expression in LS174T cells. H-I. Analysis of the cytotoxicity of the CS prepared from the NF-κB suppressive strains in LS174T (Panel H) and Caco-2 (Panel I) reporter cells. CS prepared from these strains did not exhibit cytotoxic effects (mean (SD)). * *p*<0.05, ** *p*<0.01, *** *p*<0.001, **** *p*<0.0001 as determined by one-way ANOVA with Dunnett’s multiple comparison test.

**Supplementary Figure 2.**
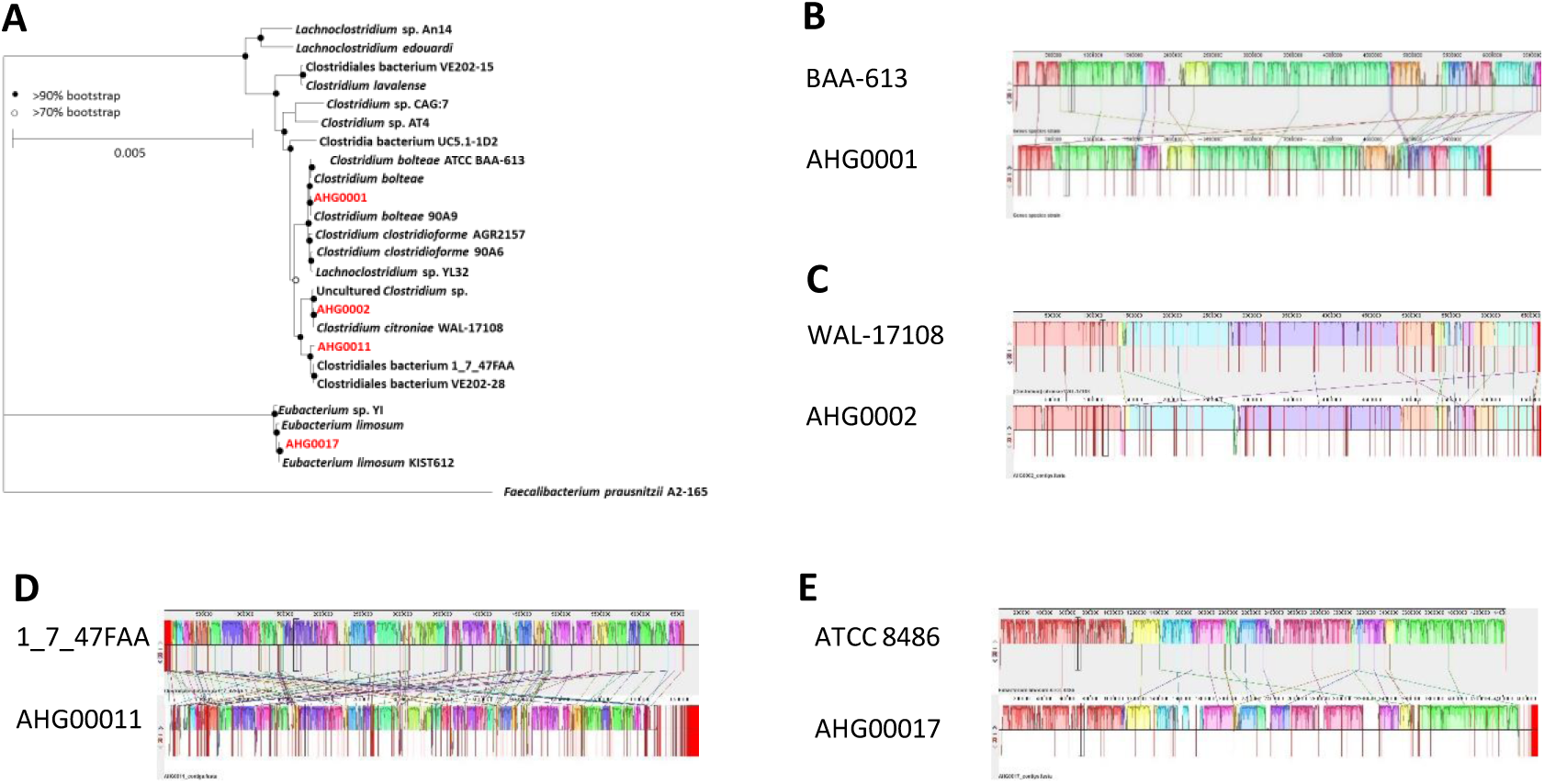
**A.** GTDB-based phylogeny of *C. bolteae* AHG0001, *C. citroniae* AHG0002, *C. aldenense* AHG0011 and *E. limosum* AHG0017 (red typeface) as determined from the concatenation of 120 universal bacterial-specific marker genes. Representative strains are included for comparative purposes (black typeface). The bootstrap values are indicated using a cut-off of >70 or >90%. B-E. The extent of genome synteny between *C. bolteae* ATCC BAA-613 and *C. bolteae* AHG0001 (Panel B), *C. citroniae* WAL-17108 and *C. citroniae* AHG0002 (Panel C), *C. aldenense* 1_7_47FAA and *C. aldenense* AHG0011, and; *E. limosum* ATCC8486 and *E. limosum* AHG0017. The red lines indicate the boundaries of chromosomes, plasmids or contigs.

**Supplementary Figure 3.**
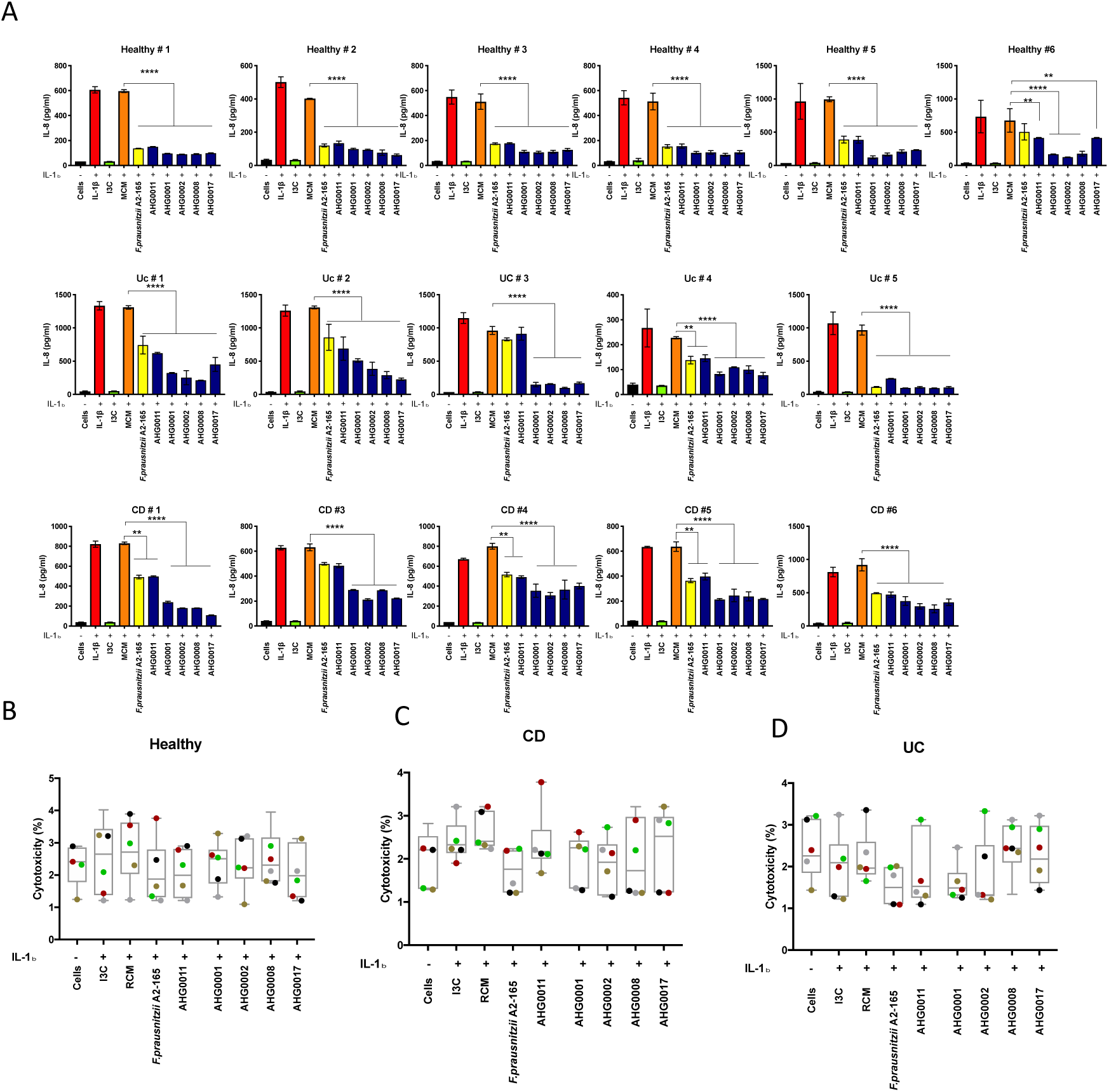
**A.** Analysis of the ability of CS prepared from the NF-κB suppressive strains to suppress IL-8 secretion in organoids produced from healthy (Healthy, n=6), Crohn’s disease (CD, n=5) or ulcerative colitis (UC, n=5) subjects. B. Analysis of the cytotoxicity of the CS prepared from the NF-κB suppressive strains in organoids produced from healthy (Healthy, n=6), Crohn’s disease (CD, n=5) or ulcerative colitis (UC, n=5) subjects. ** *p*<0.01, **** *p*<0.0001 as determined by one-way ANOVA with Dunnett’s multiple comparison test.

**Supplementary Figure 4.**
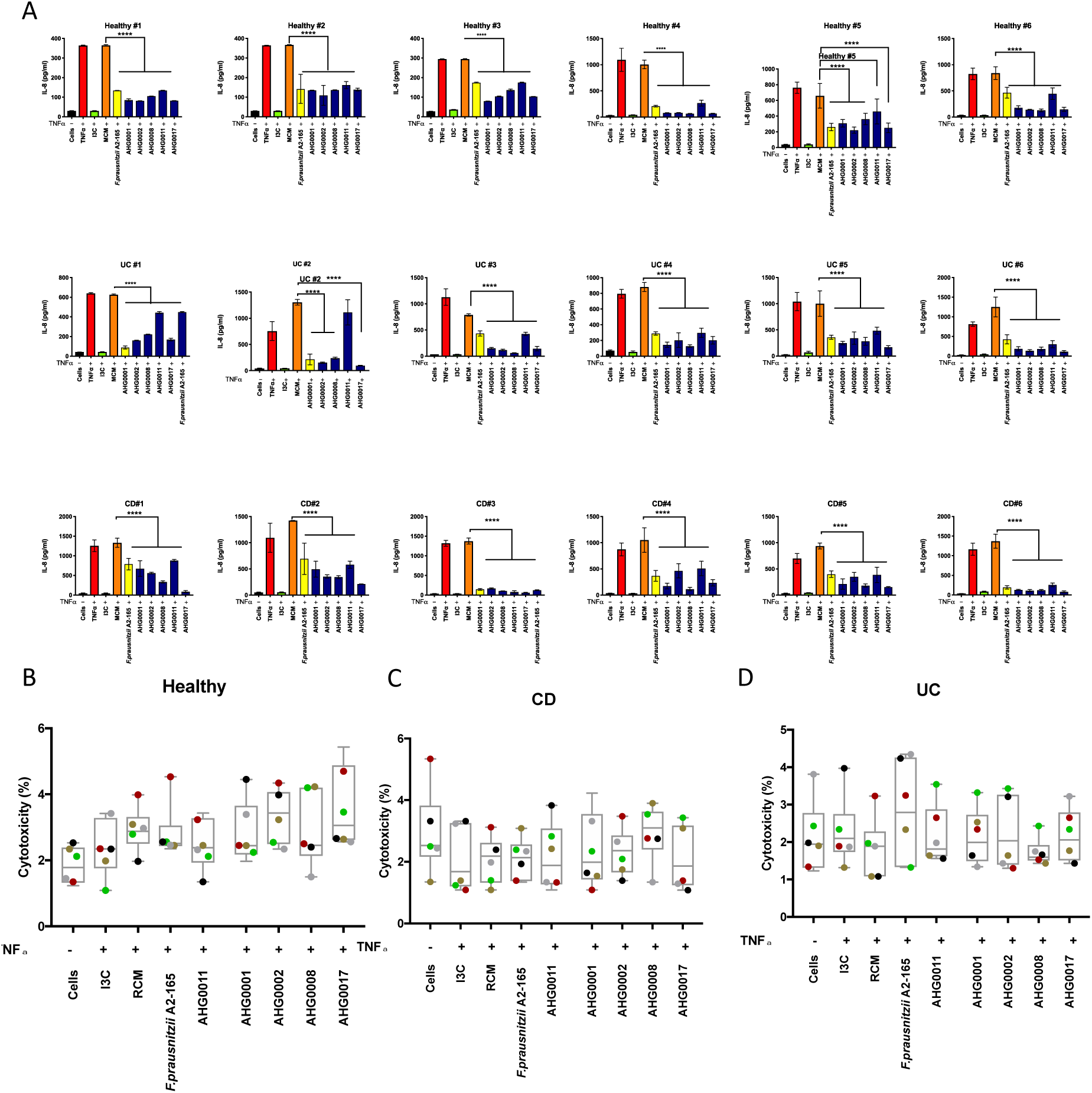
Analysis of the ability of CS prepared from the NF-κB suppressive strains to suppress IL-8 secretion in PBMCs prepared from healthy (Healthy, n=6), Crohn’s disease (CD, n=6) or ulcerative colitis (UC, n=6) subjects. **B.** Analysis of the cytotoxicity of the CS prepared from the NF-κB suppressive strains in PBMCs produced from healthy (Healthy, n=6), Crohn’s disease (CD, n=5) or ulcerative colitis (UC, n=5) subjects. * *p*<0.05, **** *p*<0.0001 as determined by one-way ANOVA with Dunnett’s multiple comparison test.

**Supplementary Figure 5.**
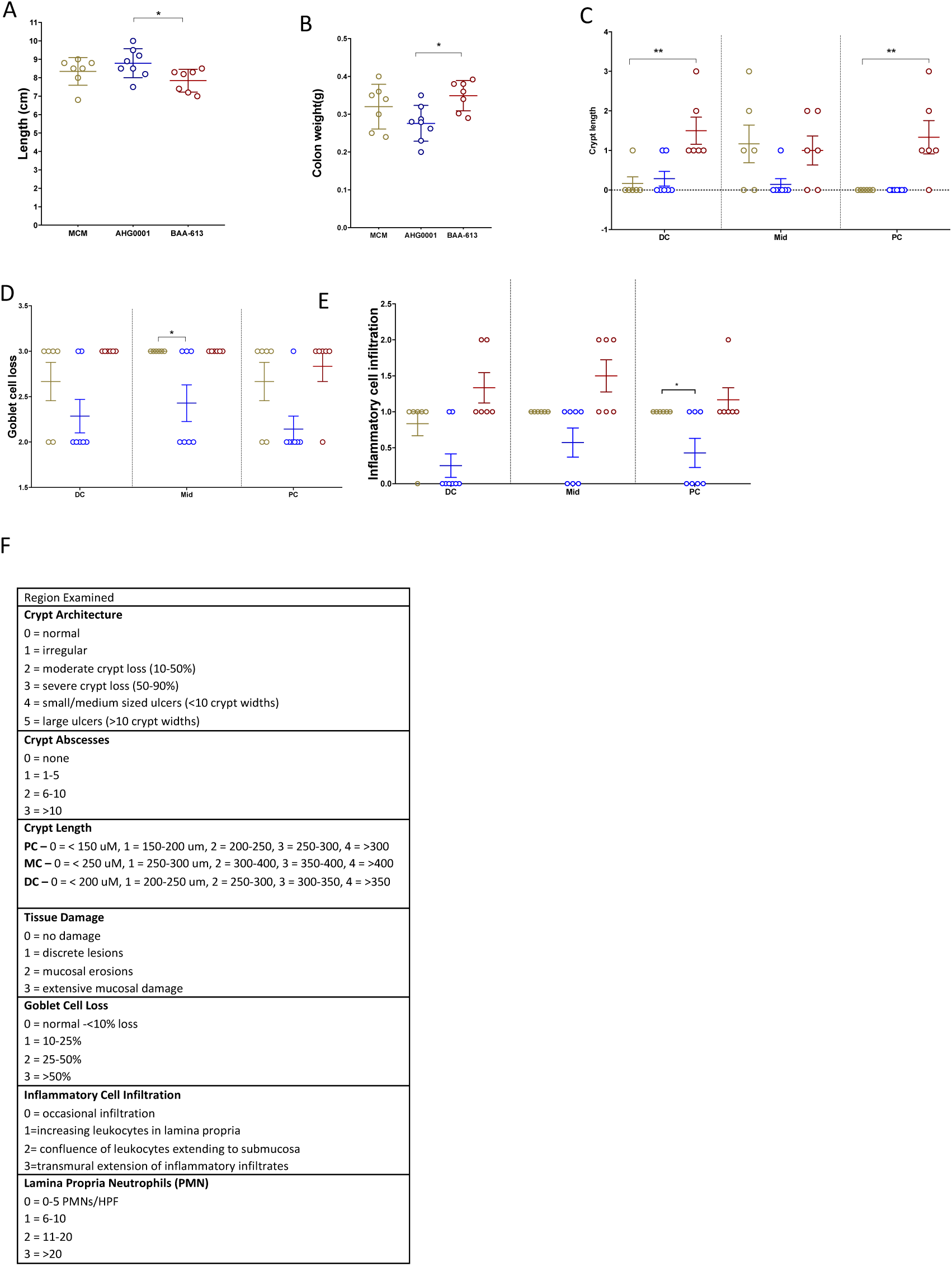
Histological colonic inflammation sub-scores following the treatment with MCM, *C. bolteae* AHG0001 and *C. bolteae* BAA-613. A. Colon length B. Colon weight C. Crypt length D. Goblet cell loss E. Inflammatory cell infiltration F. Criteria for histology sub score \**p*<0.05, ** *p*<0.01 as determined by one-way ANOVA with Dunnett’s multiple comparison test.

**Supplementary Figure 6.**
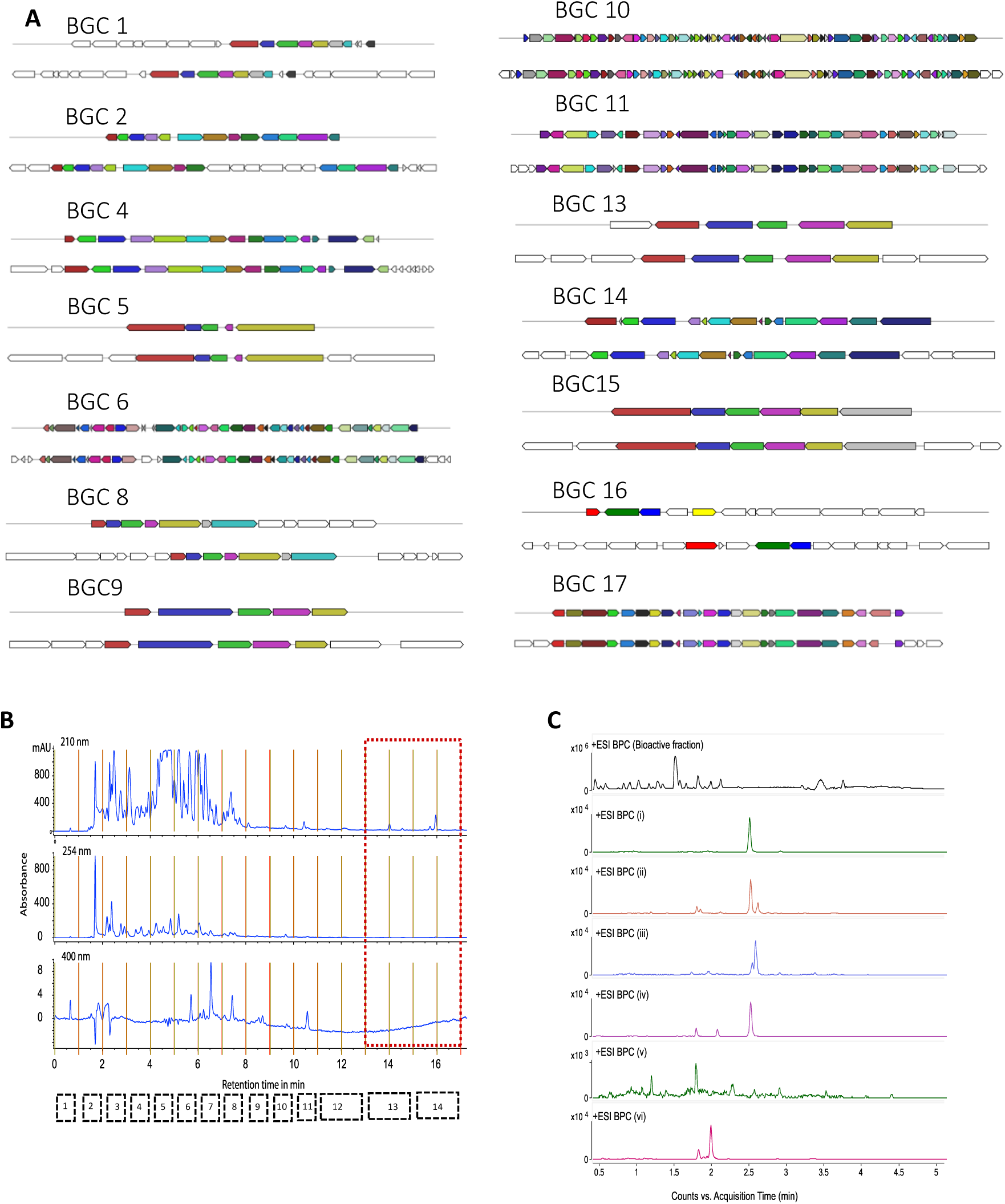
**A.** Gene organisation of the highly and partially conserved BGCs between *C. bolteae* AHG0001 (top) and *C. bolteae* ATCC BAA-613 (bottom). BGC 12 and BGC 13 are contiguous in *C. bolteae* AHG0001 and this BGC disrupted by a transposon insertion in *C. bolteae* ATCC BAA-613. **B.** Semi-preparative HPLC fractionation of the EtOAc extract of *C. bolteae* AHG0001 cultured in MCM media. The collected fractions are indicated with black dashed boxes and the red dashed box represents the NF-κB suppressive fractions. **C.** UPLC-QTOF single ion extraction chromatograms demonstrating that small molecules i-vi (Figure 5Ca) are present in the combined NF-κB suppressive semi-preparative HPLC fractions (see Panel B). BPC = selected base peak (*m/*z molecular ion) chromatogram. Small molecules vii-xvi were not present in the active fraction.

**Supplementary Table 1.**

| **Details** | **Organoids** |  |  | **PBMCs** |  |  |
| --- | --- | --- | --- | --- | --- | --- |
|  | **Healthy** | **CD** | **UC** | **Healthy** | **CD** | **UC** |
| Age average±SD | 44±12 | 40±10.71 | 38±14.81 | 25±6.39 | 47±19.57 | 31±11.89 |
| (range) | (29-60) | (25-54) | (23-57) | (21-35) | (24-74) | (21-52) |
| Sex | 3Male | 1 Male | 4 Male | 2 Male | 3 Male | 4 Male |
|  | 3Female | 4 Female | 1 Female | 4 Female | 3 Female | 2 Female |
| Severity | N/A | 2 none | 1 none | N/A | 2 none | 3 none |
|  |  | 3 mild | 1 mild |  | 4 moderate | 1 mild, |
|  |  |  | 3 moderate |  |  | 2 moderate |
| Medication | none | 1 anti-TNF, | 2 IM, | none | 5 IM, | 2 IM, |
|  |  | 1 anti-TNF+IM, | 2 IM+ 5-ASA, |  | 1 anti-TNF | 1 5-ASA, |
|  |  | 1 IM, | 1 none |  |  | 1 5-ASA+IM, |
|  |  | 1 5-ASA, |  |  |  | 1 IM +anti-TNF, |
|  |  | 1 none |  |  |  | 1 none |

IM = Immunomodulator, 5-ASA = 5-aminosalicylic acid

**Supplementary Table 2.**
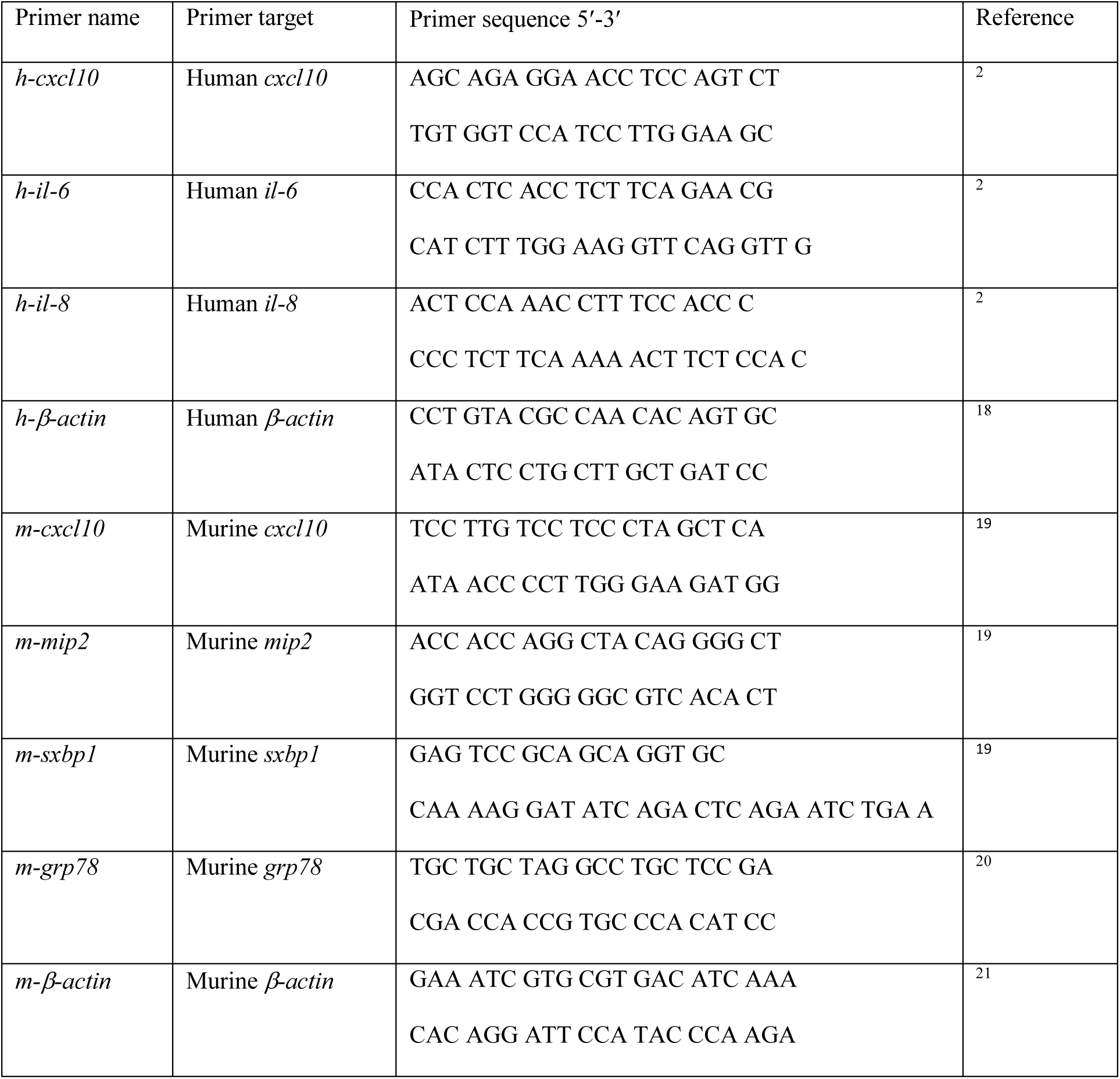

